# Human bladder organoids model urinary tract infection and bacteriophage therapy

**DOI:** 10.1101/2025.07.30.667685

**Authors:** Jacob J. Zulk, Clare M. Robertson, Samantha Ottinger, Amal Kambal, Alejandra Rivera Tostado, Rachel C. Fleck, Allyson E. Shea, Cristian Coarfa, Sarah E. Blutt, Anthony W. Maresso, Kathryn A. Patras

**Affiliations:** Department of Molecular Virology and Microbiology, Baylor College of Medicine, Houston, TX, USA; Tailored Antibacterials and Innovative Laboratories for Phage (Φ) Research (TAILΦR), Baylor College of Medicine; Department of Microbiology and Immunology, University of South Alabama, Mobile, AL, USA; Department of Molecular and Cellular Biology, Baylor College of Medicine.; Alkek Center for Metagenomics and Microbiome Research, Baylor College of Medicine.

## Abstract

Urinary tract infections (UTIs), primarily caused by uropathogenic *Escherichia coli* (UPEC), are among the most common antibiotic-resistant infections. Despite this, currently available preclinical UTI models lack the breadth of morphotypic and heterogenous cell populations of the human bladder, impairing the development of novel therapies. To address these limitations, we developed human bladder organoids derived from the bladder stem cells of multiple healthy donors which recapitulate cellular diversity of the urothelium. Using bulk and single cell RNA-sequencing, we characterized organoid responses to UPEC and phage exposure individually and in combination to model phage therapy. Although phage minimally affected the uroepithelium in the absence of infection, during UTI, phage treatment reduced bacterial burdens and dampened inflammatory responses and barrier disruption. Collectively, our findings highlight human bladder organoids as a tool for capturing conserved and individual-specific uroepithelial responses to infection while also providing preclinical efficacy and safety testing for therapeutic development.

## INTRODUCTION

Urinary tract infections (UTIs) are among the most prevalent bacterial infections globally and are predominantly caused by uropathogenic *Escherichia coli* (UPEC)^1–4^. UTIs represent the second most common reason for antibiotic prescriptions in U.S. adult outpatient care^5^; however, UTI recurrence rates remain high (20-30%) and repeated antibiotic use remains the standard of care^6,7^. Alarmingly, antimicrobial-resistant (AMR) uropathogens were associated with or attributable to nearly 300,000 deaths worldwide in 2019 alone^8^, underscoring an urgent need for alternative therapies to combat this growing public health threat.

Both *in vivo* and *in vitro* models are extensively used to investigate UTI pathogenesis, host responses, and therapeutic efficacy^9,10^. *In vivo* models are particularly valuable for studying long-term infection dynamics, immune responses, and health outcomes; yet physiological and immunological differences between animal models and humans limit the direct translation of findings^9–13^. Conversely, *in vitro* cell monolayer models facilitate the study of human bladder epithelial (uroepithelial) responses and can be adapted to model urine voiding dynamics^14–16^. These models, however, lack the complex multilayered architecture of the human bladder. To bridge this gap, three-dimensional bladder organoid models offer a physiologically relevant system that more closely resembles the morphology and heterogeneity of the human uroepithelium. While many of these models were developed to study bladder cancers^17–21^, several have been generated with UTI research in mind^22–26^. A few organoid models have incorporated human urine, with reported impacts on both human and bacterial cells^23,27,28^. As a critical limitation, the cells used for these models are rarely derived from healthy reproductive-aged women, a key demographic in studying UTI and an important consideration given sex-based differences in response to UTI^29^.

Bacteriophage (phage) therapy poses a promising tool to circumvent rising AMR in uropathogens. Phages are viruses that specifically infect bacteria, and their potential therapeutic activity, including for UTIs, has been recognized for a century^30–33^. Although interest in phage therapy declined with widespread implementation of antibiotics, the growing threat of AMR and concerns over antibiotics’ unintended effects on the microbiome^34^ have reignited interest in phage-based treatments. While phage therapy is generally regarded as safe, phage therapy can elicit both innate and adaptive immune responses^35–43^. Furthermore, mounting evidence supports phages can bind to and be internalized by multiple human cell types^44–53^. Despite improvements in human bladder modeling and renewed interest in phage therapy, bladder organoid models have not yet been implemented in the context of phage therapy development for UTI. Such studies could yield critical insights into the safety and efficacy of phage treatment in a more physiologically relevant system.

Here, we developed human bladder stem cell-derived organoids from multiple healthy donors that recapitulate hallmark features of the bladder uroepithelium including transcriptionally distinctive cell types. Using these organoids, we evaluated uroepithelial responses to UPEC infection or phage exposure individually and in combination as a model for phage therapy during UTI in humans. Although phage minimally affected the uroepithelium in the absence of infection, during UTI, phage treatment initially reduced bacterial burdens and dampened host inflammatory responses to UPEC with diminished effects at later timepoints. Collectively, our findings highlight human bladder organoids as a tool for capturing conserved and individual-specific uroepithelial responses to bacterial pathogens while also providing valuable preclinical efficacy and safety testing of novel therapeutics in a human-relevant system.

## RESULTS

### Human bladder organoids recapitulate key features of the human urothelium

Human bladder organoids (HBOs) were generated from primary bladder dome epithelial cells obtained from 2 male and 2 female donors aged 18–30 to establish four genetically distinct HBO lines (**Fig. 1A-B**). To determine similarities between HBOs and human bladder tissue *in situ*, we performed bulk RNA sequencing of HBO lines and compared normalized transcripts of 87 bladder-associated protein-coding genes^54^ with human bladder reference tissue from the Human Protein Atlas^55,56^. Although expression levels varied across the four HBO lines, all lines clustered more closely with bladder reference tissue compared to other human tissue types (**Fig. 1C**) (**Table S1**).

**Figure 1:**
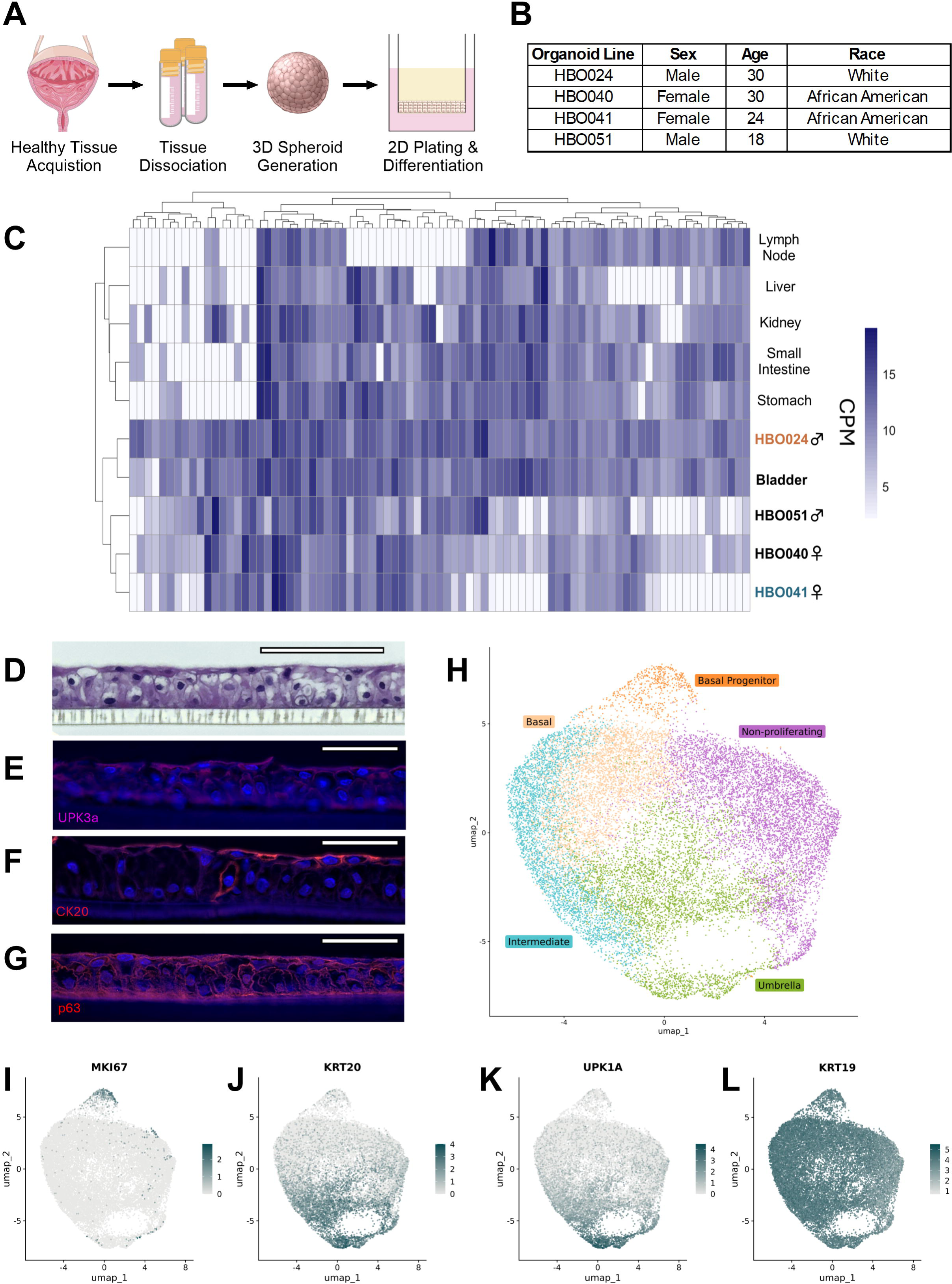
Human bladder organoids (HBOs) recapitulate key features of the bladder. (**A**) Schematic describing the generation of the HBO model. (**B**) Demographic information from bladder cells used to generate HBOs. (**C**) Hierarchical clustering of normalized counts per million (CPM) of the four HBO lines (HBO024, 040, 041, and 051) and human reference tissue in a subset of 87 bladder-associated genes. (**D**) H&E microscopy of HBO041. Scale bars represent a length of 110 µm. Immunofluorescence microscopy of HBO041, staining for (**E**) UPK3a, (**F**) CK20, and (**G**) p63. Scale bars for immunofluorescence represent a length of 50 µm. (**H**) UMAP projection of HBO041 organoids with cell populations identified. Projections were used to evaluate expression of urothelial markers including (**I**) MKI67 (Ki-67), (**J**) KRT20, (**K**) UPK1A, and (**L**) KRT19. (C) Two or (H-L) three organoid transwells for each line were independently grown and cells were pooled prior to RNA extraction and sequencing. Images used in (A) are provided by Servier Medical Art (https://smart.servier.com/), licensed under CC BY 4.0, by Free SVG, licensed under CC0, or by NIAID NIH BIOART Source (bioart.niaid.nih.gov/bioart/88 and bioart.niaid.nih.gov/bioart/399).

The urinary bladder urothelium is composed of multiple cell populations including basal, intermediate, and umbrella (apical) cells which have unique properties and functions^57–60^. We observed the formation of multilayered cell organization when differentiated on transwells, with elongated and flattened umbrella cells present on the apical surface (**Fig. 1D**), resembling key features of native bladder uroepithelium. Likewise, using immunofluorescent staining, we observed apical expression of uroplakin-3A and cytokeratin-20, consistent with observations of the urothelium as well as of other bladder organoid models^22–24,26,61^ (**Fig. 1E-F**). Important for squamous epithelial proliferation and differentiation^62^, we observed p63 expression throughout the organoid layer (**Fig. 1G**).

To further characterize the cellular landscape of HBOs, we performed single-cell RNA sequencing of differentiated organoid line HBO041. Unsupervised clustering, UMAP dimensional reduction, and marker identification identified five distinct clusters representing cell types corresponding to non-proliferating, basal, basal progenitor, intermediate, and umbrella cells (**Fig. 1H**) (**Table S1**). Cell cycle marker MKI67 (Ki-67) expression was concentrated in the basal progenitor cell population (**Fig. 1I**), whereas cytokeratin-20 (KRT20) and uroplakin-1A were primarily expressed in the umbrella cell cluster (**Fig. 1J-K**). These are contrasted with cytokeratin-19 (KRT19), which is expressed throughout the urothelium and throughout HBO041^61^ (**Fig. 1L**).

### Uropathogenic *E. coli* (UPEC) colonizes and invades HBOs resulting in an inflammatory response and barrier disruption

We selected two HBO lines, HBO024 and HBO041, for further study based on their distinct gene expression profiles (**Fig. 1C**) and their derivation from male (HBO024) and female (HBO041) donors. During infection, UPEC directly adheres to the umbrella cells, invades via a zipper mechanism, and establishes intracellular reservoirs^63–67^. To model UPEC infection in HBOs, we removed the apical media from HBO transwells and replaced it with pooled human urine containing 10LJ colony-forming units (CFU) of UPEC strain UTI89. After two hours of infection, approximately 10LJ CFU of bacteria were adhered to HBOs, and this increased to 10^6–7^ CFU per transwell after an additional two hours (**Fig. 2A**). Bacterial levels were comparable between the HBO024 and HBO041 lines at both time points.

**Figure 2:**
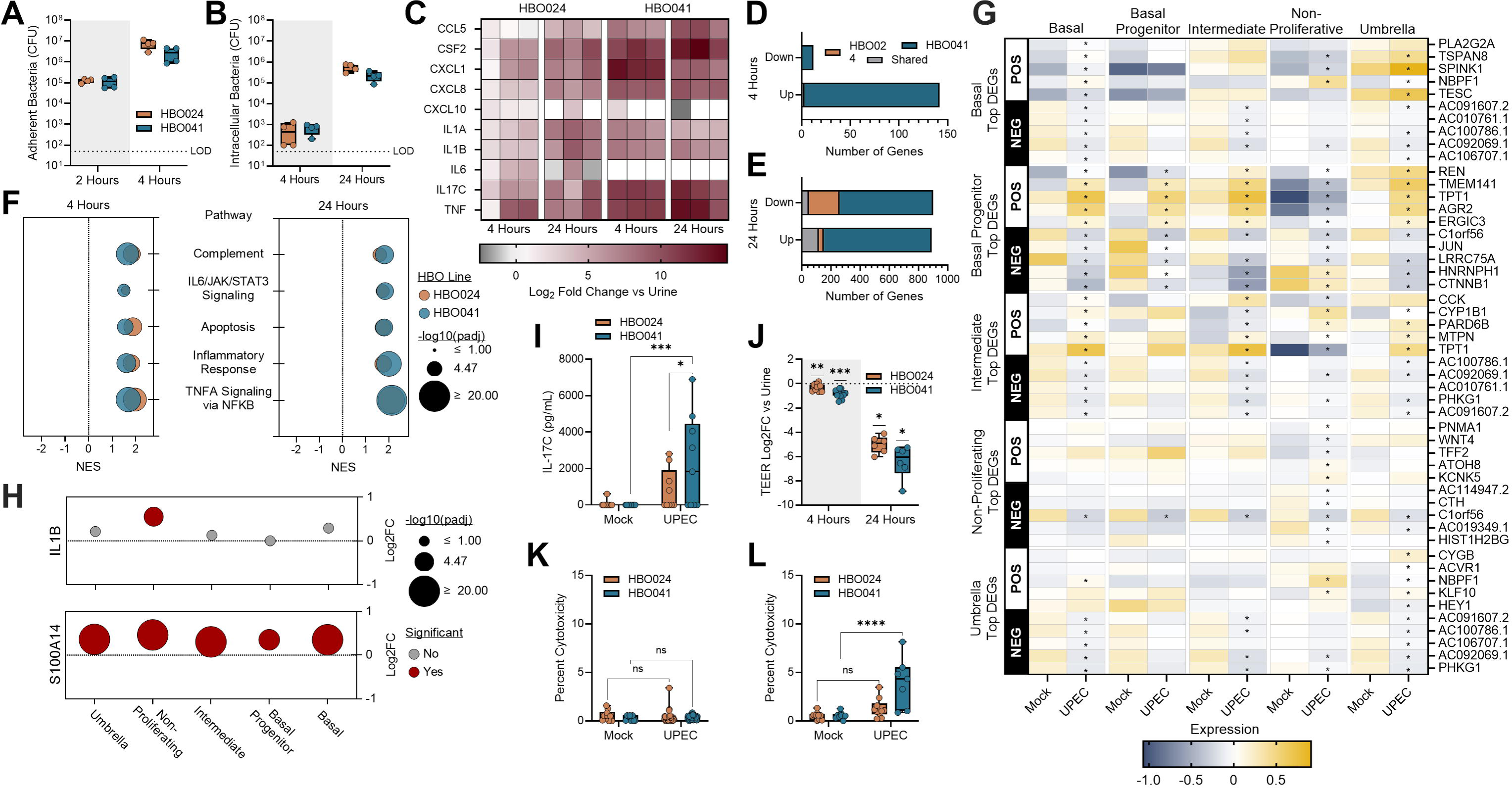
HBOs model multiple aspects of urinary tract infection. (**A**) Adherent bacteria and (**B**) intracellular bacteria recovered at 4- and 24-hours post-infection. Bulk RNA-sequencing of infected HBOs to (**C**) quantify cytokine expression changes and (**D, E**) counts of the overall number of DEGs. (**F**) Hallmark pathway analyses at 4- and 24-hours post-infection. (**G**) Single-cell RNA-sequencing was used to generate a heatmap showing the five most up- and down-regulated genes from each cluster after UPEC infection and the change in RNA expression of those genes amongst the other cell clusters. (**H**) Single cell RNA-sequencing was also used to quantify the induction of *IL1B* and *S100A14* following UPEC infection for each cell cluster. (**I**) IL-17C production in apical urine at 24-hours post-infection. (**J**) Trans-epithelial electrical resistance (TEER) across the transwell insert at 4- and 24-hours post-infection. Lactate dehydrogenase release in apical urine at 4- (**K**) and 24-hours (**L**) post-infection. Box and whisker plots show median and min/max with points representing individual organoid transwells. Data is representative of 2 (A-B), 3 (C-F), or 4+ (J-L) independent experiments. Statistical comparisons between control and UPEC infected groups were made using Kruskal-Wallis tests with Dunn’s multiple comparisons (A-B), Wilcoxon Rank Sum with Bonferroni correction (G-H), or two-way ANOVA tests with uncorrected Fisher’s LSD (I, K-L). Generalized linear model with Log2 fold change >1 and Wald tests with FDR adjusted p-valueLJ<0.05 was used to determine DEGs (D-E). Wilcoxon signed-rank test was used for testing the effect of infection on TEER, relative to urine-treated controls (J). Gene set enrichment analyses were conducted using the hallmark gene set collection from the Molecular Signatures Database (F,H). *p<0.05, **p<0.01, ***p<0.001, ****p<0.0001.

UPEC invasion of the uroepithelium enables evasion of host immune defenses and antibiotic treatment^64,68^. Using a gentamicin protection assay, we detected intracellular UPEC at 4 hours post-infection, with bacterial counts ranging from 10² - 10³ CFU per transwell (**Fig. 2B**). When gentamicin exposure was extended for an additional 20 hours, intracellular bacterial counts reached 10LJ - 10LJ CFU, indicating intracellular replication.

To examine the host response to UPEC infection, HBOs were inoculated with 10LJ CFU of UPEC and incubated for 4 to 24 hours. Following infection, apical media was collected for cytokine and cytotoxicity measurements, and cells were collected for bulk RNA sequencing. Compared to treatment with urine alone, infected HBOs upregulated multiple inflammatory genes commonly associated with UTI response at both time points including *CXCL8* (*IL8*), *CCL5*, *IL1B*, and *TNF*^69–73^ (**Fig. 2C**, **Fig. S1A-B**). Interestingly, *IL17C*, whose role in UTI response is undefined^74,75^, showed increased gene expression in both HBO lines and timepoints consistent with other infection response genes. Other UTI response genes, such as *IL6*^76,77^ and *CXCL10*^78^, were inconsistently impacted by infection across HBO lines. At 4 hours post-infection, the two HBO lines showed minimal overlap in differentially expressed genes (DEGs) with HBO041 displaying a more robust acute UPEC response compared to HBO024 (**Fig. 2D, Table S2**). At 24 hours post-infection, the total number of DEGs increased in both lines, but again, HBO041 displayed a more extensive response, with minimal overlap to HBO024 (**Fig. 2E**).

While the overall number of DEGs varied between the two HBO lines, pathway analyses revealed considerable congruency. In both lines, pathways associated with the host response to infection, including TNFα signaling via NF-κB, IL6/JAK/STAT3 signaling, and apoptosis, were significantly upregulated within 4 hours post-infection and remained elevated at 24 hours (**Fig.** 2F, Table S3).

To characterize UPEC responses within individual cell populations, we conducted single-cell sequencing on HBO041 infected with UPEC for four hours. We retained antibiotics in the basal HBO media to minimize the impact of cell death (apoptosis) pathway activation. We identified the top five up- and down-regulated genes for each of the five cell clusters (**Fig. 2G**) and compared their changes in expression across all clusters. While many of the DEGs have unknown associations with infection, some differentially regulated genes are associated with cell proliferation and infection. NBPF1, which negatively regulates cell proliferation^79^, was increased in basal, non-proliferating, and umbrella cell populations following infection. AGR2 was among the most upregulated genes following infection in the basal progenitor cluster. AGR2 ensures that newly synthesized proteins are properly folded, and its production is associated *Enterobacteriaceae* abundance in children with Crohn’s disease^80,81^. AGR2 was significantly upregulated following infection in all other cell clusters except for the non-proliferative cluster, where it is significantly downregulated following infection. We also examined markers of immune modulation within cell clusters following infection. UPEC infection led to a statistically significant increase in *IL1B* transcription in the non-proliferating cell population, whereas infection significantly increased transcription of *S100A14*, a member of the S100 protein family, in all cell clusters (**Fig. 2H**).

Although minimally described in response to UTI, IL-17C is an epithelial-derived cytokine produced in response to proinflammatory cytokines including IL-1β and TNF, tissue damage, or microbial infection^74,82–84^. We observed increased IL-17C production via ELISA at the 24-hour timepoint in both HBO lines (**Fig. 2I**). To determine whether IL-17C could be detected in clinical samples, we measured IL-17C via ELISA in urine samples from female patients with UTI, asymptomatic bacteriuria, or healthy controls. Detection of IL-17C was rare (4 of 28 total samples) and levels were not different between the three groups (**Fig. S1C**).

During UTI, host responses trigger the exfoliation of apical umbrella cells through apoptosis-like mechanisms^67,68,85^ and decrease tight junction expression resulting in compromised urothelial integrity^77,86,87^. To assess whether our HBO model recapitulates these features, we measured transepithelial electrical resistance (TEER) in infected transwells. A decline in HBO barrier integrity, measured by decreased TEER values, was evident within 4 hours of infection, with further disruption of barrier integrity observed by 24 hours (**Fig. 2J**). This change in barrier integrity at 4 hours post-infection was not correlated with a change in expression of tight junction proteins (**Fig. S2A**). After 24 hours of infection, minor changes in expression were noted, including an increased expression of claudin-4 (**Fig. S2B, S2C**), which is present in the tight junctions of umbrella cells^88^. Despite loss of electrical resistance, no significant differences in cytotoxicity were measured at 4 hours post-infection (**Fig. 2K**), and only minor increases in cytotoxicity compared to urine were observed at 24 hours post-infection, reaching only ∼1% of HBO024 and ∼4.5% of HBO041 cells at 24 hours post-infection (**Fig. 2L**).

### Phage exposure minimally impacts HBO transcription and barrier function

When used intravenously, bacteriophages stimulate both innate and adaptive immune responses^89^. Recent studies suggest that phages applied directly to the bladder may be comparatively less immunogenic^90^. To assess uroepithelial responses to phage in the absence of infection, we treated HBOs with pooled human urine containing 10^8^ PFU of a well-characterized, lytic, and clinically-applied phage, HP3^44,91–94^, for 4 or 24 hours. In the HBO024 line, very few DEGs were identified compared to urine alone controls at both 4 (**Fig. 3A**, **Table S4**) and 24 hours (**Fig. 3B**), and no DEGs were conserved between the two timepoints.

**Figure 3:**
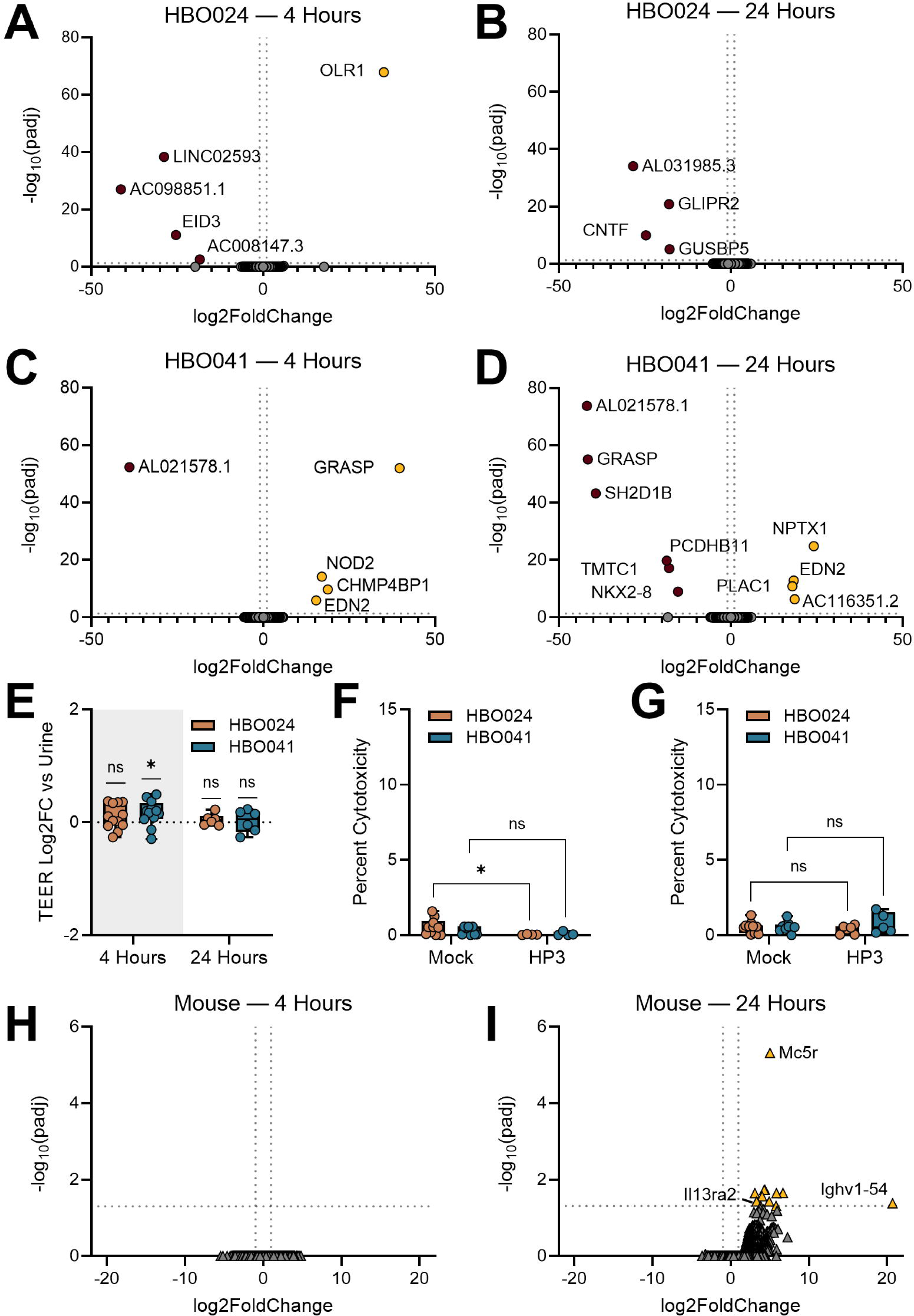
Bacteriophage HP3 minimally alters HBO and murine bladder transcriptional profiles. Differentially expressed genes for HBO024 (A-B) and HBO041 (C-D) after 4- and 24 hours of incubation with phage HP3. (E) Trans-epithelial electrical resistance (TEER) across the transwell insert at 4- and 24-hours post-treatment. Lactate dehydrogenase release in apical urine at 4- (F) and 24-hours (G) post-treatment. Male and female mice were treated with 10^8^ PFU of HP3 and bladders were collected at (H) 4 or (I) 24 hours for bulk RNA sequencing and DEG analyses. Volcano plots were generated from 3 independent replicates of each organoid line (A-D) or (H-I) from a combined 8 HP3 treated and 4 mock treated bladders at each timepoint. Generalized linear model with Log2 fold change >1 and Wald tests with FDR adjusted p-valueLJ<0.05 was used to determine DEGs (A-D, H-I). (E) TEER and (F-G) cytotoxicity measurements were made over 4-6 independent experiments. Box and whisker plots show median and min/max with points representing individual organoid transwells. Wilcoxon signed-rank test was used for testing the effect of infection on TEER, relative to urine-treated controls (E). Two-way ANOVAs with uncorrected Fisher’s LSD test were used to measure differences between phage treated and untreated HBOs (F-G). *p<0.05.

Likewise, HBO041 had few DEGs in response to phage treatment at 4 (**Fig. 3C**) and 24 hours (**Fig. 3D**). Among these, upregulated *EDN2*, encoding endothelin 2, and downregulated AL021578.1, encoding a pyroptosis-related lncRNA^95^, were observed at both timepoints. Of note, *NOD2* expression, encoding the intracellular innate immune sensor NOD2^96^, was increased after 4 hours of exposure but not at the 24-hour timepoint. Under these same conditions, TEER of phage-treated HBOs remained comparable to that of urine alone across lines and timepoints (**Fig. 3E**) and no significant change in the expression of tight junction proteins was observed (**Fig. S2D-F**). Overall cytotoxicity was low (<2%) and few differences were observed between phage treatment and urine alone at either time point (**Fig. 3F-G**).

To compare HBO responses with whole bladders *in vivo*, we transurethrally inoculated wild type C57BL/6J male and female mice with HP3 or a mock phage preparation. Murine bladders were collected at 4 and 24 hours post-treatment and subjected to bulk RNA-sequencing. In line with our *in vitro* findings, we detected minimal transcriptional responses to phage treatment. No DEGs were detected at 4 hours (**Fig. 3H**) but several genes were upregulated in response to phage treatment at 24 hours (**Fig. 3I, Table S4**). These included *Mc5r*, related to immune response and regulation^97^ and *Ighv1-54* (immunoglobulin heavy variable V1-54), predicted to be involved in immunoglobulin-mediated immune responses. In addition to the murine model of phage treatment, we compared phage responses between HBOs and a widely used immortalized bladder epithelial cell line HTB-9 (5637). No DEGs were observed in HTB-9 cells treated with phage HP3 at either 4 or 24 hours of incubation (**Table S4**), although statistical power was limited by the number of replicates in these experiments. Taken together, these results suggest that HBOs, as well as widely adopted *in vivo* and cell culture models, do not mount robust innate immune responses to the bacteriophage HP3.

### Bacteriophage HP3 reduces adherent bacteria and dampens initial host responses but fails to eliminate UPEC infection

To test the therapeutic potential for HP3 to limit UPEC infection of the uroepithelium, HBOs were infected with UPEC for two hours, then apical urine was replaced with urine alone or urine containing 10^8^ PFU of HP3. Two hours after phage treatment, adherent bacterial counts were reduced by 94% and 99% for HBO024 and HBO041, respectively (**Fig. 4A**). As infection progresses, UPEC invade uroepithelial cells, evading host immune responses and antibiotic therapy^64,65,98–101^. To test the impact of phage on UPEC intracellular reservoirs, HBOs were infected with UPEC in urine for two hours followed by replacement with urine containing gentamicin or gentamicin and HP3. After one hour, urine was replaced with a lower maintenance dose of gentamicin (10 µg/mL) with or without HP3. In contrast to HP3-mediated reduction of extracellular UPEC, there was no effect of HP3 treatment on intracellular bacteria at 24 hours after UPEC infection in either HBO line (**Fig. 4B**).

**Figure 4:**
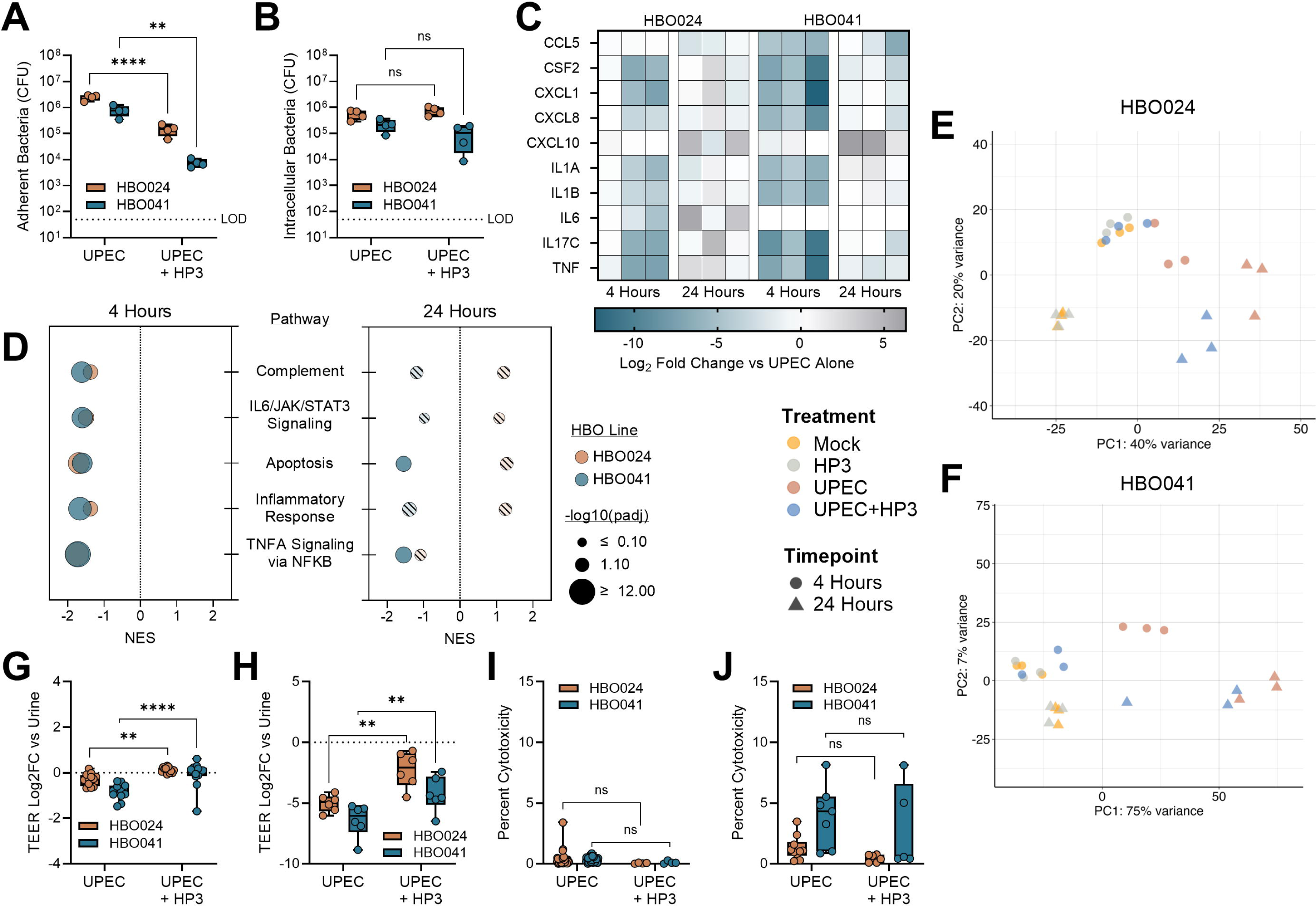
Phage treatment reduces extracellular UPEC and HBO inflammatory responses early in infection. Phage HP3 was tested for its ability to remove established (A) extracellular bacteria and (B) intracellular bacterial reservoirs. Bulk RNA-sequencing of infected HBOs with or without HP3 treatment was used to quantify (C) cytokine expression changes and (D) to generate pathway analyses at 4- and 24-hours post-infection. Solid colored circles in pathway analysis represent significant results while crosshatched circles represent non-statistically significant findings. Principal component analyses of (E) HBO024 and (F) HBO041 that were untreated, treated with HP3, treated with UPEC alone, or with UPEC and HP3. Changes in HBO trans-epithelial electrical resistance (TEER) across the transwell insert at (G) 4- and (H) 24-hours post-infection and compared between phage and no phage groups. Lactate dehydrogenase release in apical urine at (I) 4 and (J) 24 hours. Box and whisker plots show median and min/max with points representing individual organoid transwells (A-B, G-J). Data is representative of 2 (A-B), 3 (C-F), or 4+ (G-J) independent experiments. Data were analyzed using (A-B, G-J) two-way ANOVAs with uncorrected Fisher’s LSD. Gene set enrichment analyses were conducted using the hallmark gene set collection from the Molecular Signatures Database (D). **p<0.01, ****p<0.0001.

To determine the impact of HP3 on host response to UTI, we performed bulk RNA-sequencing of HBOs infected with UPEC with or without HP3 treatment for 4 and 24 hours. Infected HBOs treated with HP3 had reduced expression of several key cytokines and chemokines at 4 hours; however, these differences were largely resolved by 24 hours (**Fig. 4C, Table S5**). Pathway analyses supported these observations, revealing that HP3 treatment decreased inflammation and apoptosis-related pathway transcription 4 hours post-infection, but this effect was mitigated at 24 hours (**Fig. 4D**, **Table S6**). Using principal component analysis, we observed that UPEC infection was the primary driver of gene expression differences between treatment groups at both 4 and 24 hours (**Fig. 4E-F**), with later timepoints extenuating group variance.

To test whether HP3 treatment altered UPEC-mediated reduction of HBO barrier integrity, we measured TEER at 4 and 24 hours. Compared to infection with UPEC alone, the addition of HP3 preserved barrier integrity at 4 hours (**Fig. 4G**) as well as 24 hours post-infection (**Fig. 4H**). However, overall cytotoxicity was comparable in the UPEC-infected HBOs and those treated with HP3 at both timepoints (**Fig. 4I-J**). These results suggest that HP3 treatment can reduce bacterial attachment and host inflammatory responses initially, but ultimately, bacterial infection can overcome phage-mediated effects, resulting in HBO immune activation and cytotoxicity.

## DISCUSSION

Urinary tract infections remain a major medical challenge due to the high prevalence and recurrence rates and increasing antibiotic resistance. These obstacles underscore the need to better understand the host–pathogen interactions during UTI and to develop alternatives to antibiotic treatment. Here, we present novel human bladder organoids derived from primary cells that closely mimic key features of the healthy human bladder uroepithelium and capture individual variations across donor lines. Our findings show that UPEC infection triggers inflammatory responses that ultimately compromise epithelial barrier function. While bacteriophages alone had minimal effects on the organoids, their presence during UPEC infection dampened early bacterial levels and inflammation, supporting their potential as a therapeutic intervention.

The urothelium is composed of basal, intermediate, and umbrella cells, with each type expressing distinct cellular markers. Using canonical cell type markers from single cell RNA-sequencing of the human bladder^59^, we classified the cells within our HBO041 organoid line into five distinct clusters: basal, intermediate, umbrella, basal progenitor, and non-proliferative cells. While phenotypically expressing markers characteristic of the differentiated urothelium, the number of HBO cell layers present was often less than the approximately seven observed in the healthy human urothelium^102^. As a key limitation, while HBOs mimic the stratified human bladder urothelium, they do not model aspects like urine flow, resident immune cells, or mechanical stretch and corresponding changes in bladder cell morphology, which are important for *in vivo* bladder physiology^9^. Additionally, only two HBO lines were used for the bulk of experiments. Future studies would benefit from expanding the number of donor lines studied, especially as sex-based and individual differences in bladder cell populations are noted in the healthy adult population^60^.

In this work, we evaluated the ability of HBOs to mirror infection phenotypes reported in other models. Both *in vitro* and *in vivo*, UPEC form intracellular reservoirs to evade antibiotics and immune detection^64,65,100,101,103^. UPEC intracellular reservoirs have been reported in murine organoids^104^, immortalized human cell line “organ-on-a-chip” models^68^, and bladder ‘assembloids’, a complex, layered assembly of urothelium, connective tissue, and muscle^24^. Consistent with these studies, we observed intracellular UPEC reservoirs in our HBOs that persisted and replicated despite antibiotic treatment. *In vivo* and in murine organoids, the bladder mounts a rapid pro-inflammatory response to infection, including production of IL-1β, IL-6, IL-8 (CXCL8), and TNF^104,105^. Our HBOs likewise increased expression of these genes within 24 hours of infection. Sharma *et al*.^104^ furthered showed that neutrophils migrated preferentially towards infected organoids, likely due to infection-induced cytokine production: a hypothesis that could be explored using transwell co-culture of HBOs with primary neutrophils.

Of particular interest, both HBO lines robustly upregulated IL-17C as early as 4 hours post UPEC infection. Unlike other IL-17 family members, IL-17C is primarily expressed by epithelial cells at mucosal surfaces in response to stimuli including live or heat-killed *E. coli*^83,84,106–108^. Prior work detected increased *IL17C* transcription in UPEC-infected bladder epithelial cells^74,75^, and here, we confirmed IL-17C production by the human uroepthelium upon UPEC infection. Once produced, IL-17C can increase expression of other cytokines and chemokines involved in pathogen clearance such as CSF3^109^, G-CSF^106^, IL-6^110^, and IL-8^110^ as well as antimicrobial peptides including human beta defensin 2 (hBD2)^106,109,110^ and lipocalin-2 (LCN2)^110^. Despite potent release of IL-17C by HBOs, we did not detect increased levels in a patient cohort of UTI. Possible explanations for this include cytokine dilution or washout below the limit of detection *in vivo* compared to the static HBO conditions or differences in cytokine induction kinetics *in vivo* versus *in vitro*. Taken together, IL-17C upregulation, with concurrent increases in other cytokines, chemokines, and antimicrobial peptides may be important for bacterial clearance and UTI resolution, but additional research is still needed.

Previous studies have shown that female mice have higher expression of inflammatory cytokines and chemokines 24 hours after UPEC infection compared to males^111^. We observed a similar trend with the female-derived line (HBO041) showing higher early inflammatory gene expression and more differentially expressed genes overall compared to the male-derived line (HBO024). Although our sample size limits statistical power, expanding the number of donor lines could enable discovery of sex-based differences in the human uroepithelium response to UPEC infection.

Consistent with previous bladder organoid models, UPEC infection disrupted HBO barrier integrity, most notably at later timepoints^22,23^. Interestingly, despite a significant loss of barrier integrity at 24 hours post-infection, lactate dehydrogenase assays indicated low cytotoxicity. This may be explained by exfoliation of live bladder cells, a known phenomenon in humans and mice with UTI^67,112–114^. Alternatively, our timepoints may have stopped short of peak cell death, despite increased expression of apoptosis-related genes. Another possibility is that UPEC-induced tight junction downregulation increased organoid paracellular permeability without causing widespread cell death^77,86^. Notably, although both HBO lines had low levels of LDH release at 24 hours, the median LDH release for HBO041 was about five times that observed in the HBO024 line, possibly reflecting genetic or sex-related differences.

To assess HBO responses to phage as a preclinical model for safety testing, we exposed HBOs to high concentrations of purified HP3 and evaluated gene expression, barrier integrity, and cytotoxicity. Phage exposure induced minimal gene expression changes and had no measurable impact on barrier function, aligning with prior studies showing little to no innate immune activation by phages^115–118^. These findings further support the well-documented safety profile of phage therapeutics^119^. Among the few differentially expressed genes, none were conserved between the two HBO lines, nor murine and HTB-9 cell comparisons. Within the HBO041 line, only two differential genes were shared at the 4 and 24 hour timepoints. One of these, *EDN2*, is a regulatory peptide involved in bladder contractility^120^. Additionally, in HBO041, *NOD2*, a bacterial and viral sensor dispensable for UTI^96,121^, was upregulated early but returned to baseline by 24 hours, suggesting a transient epithelial response to phage exposure.

During infection experiments, phage HP3 reduced UPEC burden by 90-99% within two hours, to levels at or below the original bacterial inoculum. This reduction in bacterial burden was associated with preserved barrier integrity and reduced inflammation early in infection. From our experimental design, we cannot delineate whether decreased inflammation is due to phage modulation of host responses, reduced bacterial burden, or a combination of these effects. While there were fewer live bacteria, remnants of those that were lysed, including endotoxin and other immunological triggers, were still present, and could contribute to an immune response. These observations warrant further investigation to determine the precise mechanisms underlying phage-associated immune modulation.

Previous studies have shown that bacteriophages can interact with and be internalized by mammalian cells^45–52^ with some showing intracellular phage accumulation over time. Møller-Olsen *et al*.^47^ demonstrated phagocytosis-mediated uptake of engineered K1F phage *in vitro*, leading to killing of intracellular UPEC. We tested whether phage HP3 could similarly target intracellular bacteria in HBOs, yet even after 20 hours of exposure, bacterial burdens remained unchanged. Despite the high phage input, limited interaction between the phage and organoids may have contributed to the lack of bacterial reservoir clearance. Alternatively, phages may have been taken up by cells but then either degraded by host machinery or sequestered from the intracellular bacterial compartment. Evaluating additional phages, particularly those which have been shown to interact with mammalian cells, may improve intracellular reservoir targeting.

In summary, this study is a novel integration of non-transformed bladder uroepithelial organoids with UPEC infection and phage therapy to add to the growing landscape of UTI models. Using bladder organoid lines derived from multiple individuals, we show that phage therapy can effectively disrupt initial UPEC infection while limiting inflammation and barrier disruption, further supporting phage therapeutic potential in humans. With continued development, bladder organoid models provide a powerful platform for phage screening, safety and efficacy testing, and for investigating genetic, sex-based, and pathogen-specific responses to UTI.

## MATERIALS AND METHODS

### Generation of Human Bladder Organoids (HBOs)

Primary human bladder cells were purchased from Lifeline Cell Technology (demographics in **Fig. 1B**). Frozen cell aliquots were resuspended in 90 mL of Matrigel and then 90 μL was plated into 24-well tissue culture plates and incubated at 37^°^C for 15-20 minutes. After incubation, 500 μL of HBO propagation media (DMEM-F12 + Hepes + L-Glutamax + Pen/Strep supplemented with 12.5 ng/mL FGF2, 100 ng/mL FGF10, 25 ng/mL FGF7, 5 µM A83-01, 10 µM Y-27632, 2% B27 supplement, 1.25 mM N-acetylcysteine, and 2.5 mM nicrotinamide) was added to each well and cells were returned to the incubator. Cultures were refreshed every other day with new HBO propagation media. HBOs were passaged every 6-7 days.

To passage, media surrounding the solid Matrigel was removed, washed once with PBS, fresh PBS was added and Matrigel was broken up with vigorous pipetting. Cells were transferred to a 15 mL tube and centrifuged at 400 × *g* (Eppendorf; catalog number 022626205) for 5 minutes. After centrifugation, media was removed and 0.05% Trypsin-EDTA (Gibco; catalog number 25300-054) was added to break up the cells. Tubes were then incubated at 37^°^C for 4 minutes before 500 µL of CMGF-media + 10% FBS was added. Tubes were then centrifuged at 500 × *g* for 5 minutes at 4^°^C. Cell pellets were resuspended in Matrigel at 30 µL per number of seeded wells. 30 µL of Matrigel-HBO mix was then transferred to each well of a new 24 well tissue culture plate. Matrigel was allowed to solidify for 30 minutes before HBO propagation media was added. Cultures were refreshed with new media every other day.

To generate and differentiate HBO monolayers, transwells were coated with 100 µL of collagen IV diluted in H_2_O and incubated at 37^°^C for 1.5 hours. 3-dimensional HBOs were washed with 0.5 mM EDTA and then dissociated by adding 500 µL of 0.05% trypsin/0.5 mM EDTA after removing wash media. Plates were incubated at 37^°^C for 7 minutes. To inactivate trypsin, 500 µL of CMGF-media containing 10% FBS was added to each well. Organoids were dissociated by vigorous pipetting up and down. Cell strainers were first wet with 1 mL of CMGF-media + 10% FBS then used to filter dissociated cells. Strainers were washed with an additional 1 mL of CMGF- + 10% FBS. Cells were pelleted at 400 × *g* for 5 minutes at 4^°^C then resuspended with 100 µL of HBO propagation medium for each well of the 24 well plate being harvested. After removing collagen IV from the transwells, 100 µL of cells were added to the top compartment of each. 500 µl of HBO propagation medium was added to the lower compartment of each well and plates were incubated at 37^°^C with 5% CO2 for 1 day. The following day, both apical and basal media were removed and replaced with HBO differentiation medium (DMEM-F12 + Hepes + L-Glutamax +Pen/Strep supplemented with 50 ng/mL EGF, 2% B27 supplement, and 1 mM N-acetylcysteine). Medium was then changed every other day until ready to be used for experiments. HBO lines generated reached this stage approximately 7 days after seeding.

### Bacteria and bacteriophage strains

Uropathogenic UPEC strain UTI89^103^ was used throughout this work. All bacteria were grown in LB media overnight in shaking conditions at 37^°^C prior to experiments. Phage HP3 (accession KY608967) was originally isolated from environmental sources and was prepared as previously described^91^ and stored in phage buffer^122^ at 4^°^C prior to use.

### Human urine pool preparation

Urine was collected from healthy male and female volunteers aged approximately 20 to 40 years old under approval of the Baylor College of Medicine (BCM) institutional review board (Protocol H-47537). Individual samples were pooled in equal amounts and filtered through a 0.2 µm filter (Corning; catalog number 431224) before use (*n* = 4-6 donors/pool). Filtered pooled urine was stored –80^°^C until use. New urine pools were used for each independent experiment.

### Microscopy

Organoid transwells were fixed with 4% formaldehyde for 30 minutes at room temperature before being cut out and placed in 70% ethanol prior to paraffin embedding. For immunofluorescent staining, following deparaffinization and rehydration, citrate buffer pH 6 (Genemed, catalog number 10-0020) was used for heat-mediated epitope retrieval. Slides were blocked using 2% bovine serum albumin (BSA) for two hours at room-temperature.

Primary antibodies were diluted in 2% BSA as follows and applied to samples overnight at 4^°^C: mouse anti-uroplakin-3A (1:20; Santa Cruz Biotechnology; catalog number sc166808), rabbit anti-cytokeratin 20 (1:100; Cell Signaling Technology; catalog number D9Z1Z), and rabbit anti-TP63 (1:100, Invitrogen, catalog number 703809). After washing, slides were incubated with corresponding secondary antibodies as follows for one hour at room-temperature: donkey anti-rabbit-Texas Red (1:100; Abcam; catalog number ab6800) or donkey anti-mouse-Cy5 (1:100; Invitrogen; catalog number A10524). Slides were again washed, then nuclei were stained using Hoechst (final concentration 1 µg/mL; Thermo Scientific; catalog number 62249) for 5 minutes at room temperature. Finally slides were washed and mounted using ProLong Gold antifade mountant (Invitrogen; catalog number P36934). Fluorescent slides were imaged using an Olympus IX83 inverted fluorescence microscope and processed using the Olympus cellSens Dimension imaging software (v4.1 Build 26283). H&E-stained slides were imaged using an Echo Revolve microscope (v3.2.1).

### Single-cell RNA-sequencing

After four hours of urine treatment or infection with 10^5^ CFU of UPEC in urine, cells were washed and dissociated from transwells using 0.25% trypsin (Corning; catalog number 25-053-CI) and manual dissociation in cell recovery media. Dissociated cells were filtered using a 40 µm cell strainer. Three replicates were pooled for each extraction.

Baylor College of Medicine’s Single Cell Genomics Core prepared RNA libraries using 10x Genomics kit (Chromium Single Cell 3’ v3) following their standard methods. Sequences were aligned to the human reference genome (GRCh38 2024-A pre-built reference by 10x Genomics) using the Cell Ranger pipeline (10x Genomics, v8.0.0). Subsequent analysis was performed using R (v4.3.3) Seurat (v5.1.0). Quality filtering retained cells with >200 transcripts, <4000 transcripts, and 15% mitochondrial transcripts, for a total of 19,652 cells. Data was normalized with the SCTransform algorithm, regressing out percent mitochondrial transcripts. UMAP dimensional reduction was performed, and the first 9 principal components were retained. Clusters were identified with the Leiden algorithm at a resolution of 0.3. Cell types were assigned according to each cluster’s expression of canonical bladder cell markers^59^ and each cluster’s top marker genes as determined by Wilcoxon Rank Sum test with Bonferroni correction. Non-proliferating and actively replicating progenitor clusters were confirmed via cell cycle scoring using the Seurat cell cycle gene list (2019 version). Differentially expressed genes between infected and mock-infected samples were determined for each cell type via Wilcoxon Rank Sum test with Bonferroni correction. All figures were created in R with Seurat (v5.1.0), ggplot2 (v3.5.1), or ComplexHeatmap (v2.18.0).

### Adherence and invasion assays

The day prior to the experiment, organoid transwells were switched from organoid differentiation medium to DMEM-F12 + Hepes + L-Glutamax (Gibco; catalog number 11330032) lacking antibiotics on both the apical and basolateral sides. The day of the experiment, media was removed and replaced with DMEM-F12 + Hepes + L-Glutamax basal media. Apical compartments were treated with 10^5^ CFU UPEC in 100 µL of pooled human urine. Plates were centrifuged at 120 x g for 1 minute to facilitate bacterial contact and then incubated at 37^°^C. For adherence assays, at 2- and 4-hours post infection, apical media was removed and transwells were washed three times in 1x PBS before being dislodged with 0.25% trypsin (Corning; catalog number 25-053-CI), lysed with 0.025% Triton X-100 (ChemCruz; catalog number Sc29112A), diluted, and plated on LB agar. For invasion assays, at 4 hours, apical media was removed and replaced with urine containing 100 µg/mL gentamicin for 1 hour. Transwells were then either washed with PBS, lysed, and plated, or apical media was replaced with urine containing 10 µg/mL gentamicin and incubated for an additional 19 hours.

For phage treatment assays, HBOs were infected with UPEC and incubated for two hours. After that time, apical media was removed and replaced with urine containing 10^8^ PFU of phage HP3. HBOs were incubated for another two hours before being washed and lysed as described above. For intracellular bacterial reduction assays, at two hours post infection, apical media was replaced with urine containing 100 µg/mL gentamicin with or without 10^8^ PFU of phage HP3. This was incubated for 1 hour, then apical media was replaced with urine containing 10 µg/mL gentamicin with or without 10^8^ PFU of phage HP3. At 24 hours post initial infection, HBOs were washed, lysed, and plated for CFU.

### TEER, LDH, and cytokine measurements

Trans-epithelial electrical resistance (TEER) was measured using a Millicell ERS-2 resistance system with a MERSSTX01 electrode (Millipore) at reported timepoints throughout experiments. An empty transwell was used to measure inherent transwell resistance and was subtracted from individual wells TEER measurements.

Lactate dehydrogenase (LDH) release was measured using the CyQUANT LDH Cytotoxicity Assay Kit (Invitrogen; catalog number C20301) following manufacturer directions. Samples of pooled human urine from each experiment were used to control for LDH present in the urine pool and were subtracted from the measurements. Cytokine expression was measured using human IL-17C DuoSet ELISA (R&D Systems; catalog number DY1234). Media from HBO samples was diluted 1:4 in PBS and then processed according to manufacturer’s recommendations. Human urine samples were diluted 1:2 in PBS and then processed according to the manufacturer’s recommendations.

### Human urine collection for IL-17C measurement

Human subjects research, exempt from informed consent, was approved under University of South Alabama (USA) IRB, #2178590-1, and BCM IRB, protocol H-54361. Discarded urine samples from patients with UTI, asymptomatic bacteriuria, and healthy controls were obtained from the USA Hospital Clinical Microbiology Lab and stored at –80°C.

### RNA-sequencing and analysis

Apical compartments of HBO transwells were treated with either urine alone, urine containing UPEC (10^5^ CFU), phage HP3 (10^8^ PFU), or a combination of the two (10^5^ CFU UPEC + 10^8^ PFU HP3). After 4 or 24 hours of incubation, apical and basal media were removed, transwells were carefully cut out with a scalpel, and then immediately submerged in RNAprotect Bacteria Reagent (Qiagen; catalog number 76506). Transwells in RNAprotect were frozen at –80^°^C until extraction. To extract RNA, cells were manually dislodged from the thawed samples and RNA was extracted using the RNeasy Plus Mini Kit (Qiagen; catalog number 74134).

Extracted RNA was sequenced by Novogene using the Illumina Novaseq X Plus Series platform. Mapping to reference genome GRCh38 was performed using Hisat2 (v2.0.5) and quantification was performed by featureCounts (v1.5.0-p3). Count normalization and differential gene expression analysis were performed using DEseq (v1.42.1). The packages EnhancedVolcano (v1.20.0), PCATools (v2.14.0), and pheatmap (v1.0.12) were used for data visualization in RStudio (v2024.09.0+375). Hallmark pathway analysis was performed using fGSEA (v1.28.0) with 10,000 permutations, a minimum gene set of 15 and maximum of 500. The Hallmark gene set was obtained from the Molecular Signatures Database (v2024.1.Hs) (https://www.gsea-msigdb.org/). To generate the HBO and reference tissue heatmap, consensus RNA expression data was downloaded from The Human Protein Atlas (www.proteinatlas.org, version 16). Genes for which expression was not measured in the HBO sequencing data were removed. Data were then subset by bladder elevated genes^54^ leaving a total of 87 genes which were analyzed using pheatmap in RStudio.

### Murine experiments and RNA-sequencing

All animal experiments were approved by the Baylor College of Medicine Institutional Animal Care and Use Committee (protocol AN-8233) and were performed under accepted veterinary standards. Phage treatments of male and female C57BL/6J mice (Jackson Laboratories, strain number 000664) were conducted by adapting a well-established murine UTI model described previously^123–125^. Briefly, mice were anesthetized with inhaled isoflurane and were given 10^8^ PFU of phage HP3 in PBS transurethrally via catheter with a total volume of 50 µL. Four- or 24-hours post phage instillation, bladders were removed, placed directly into RNA Protect (Qiagen; catalog number 26526) and stored at –80°C. Once all samples had been collected, tissues were disrupted in tubes containing 1.0-mm-diameter zirconia/silica beads (Biospec Products; catalog number 11079110z) using the MagNA Lyser instrument (Roche Diagnostics). RNA was then extracted using the RNeasy Plus Mini Kit (Qiagen; catalog number 74134).

Extracted murine RNA was sequenced by Novogene using the NovaSeq PE150 platform. Mapping to reference genome GRCm39 was performed, and analysis was conducted using the same methods as the HBO samples above.

### Statistical analyses

All experiments were performed with a minimum of three independent replicates except for baseline bulk and single cell RNA-sequencing (Fig. 1C, 1H-L) and bacterial colonization/killing experiments (Fig. 2A-B, Fig. 4A-B). In all figures, points represent individual organoid transwells. Data from individual experiments was combined prior to statistical analysis. Kruskal-Wallis tests with Dunn’s multiple comparisons were used throughout to compare the effects of bacteria and bacteriophage on cytokine production (Fig. 2G) and cytotoxicity (Fig. 2I-J, Fig. 3F-G), as well as differences in TEER changes in infected organoids treated with phage (Fig. 4G-H). Wilcoxon signed-rank test was used for testing the effect of bacteria and phage on TEER (Fig. 2H, Fig. 3E). Statistical analyses were performed using Graphpad Prism v10.4.2.

## ACKNOWLEDGEMENTS

This research was supported by NIH U19 grant (AI157981) to KAP and AWM. JJZ, CMR, and SEO were supported by NIH F31 training grants (DK136201, DK138748, HD111236). The Integrated Microscopy Core at Baylor College of Medicine is supported by the Center for Advanced Microscopy and Image Informatics (CAMII) with funding from NIH (DK56338, CA125123, ES030285), and CPRIT (RP150578, RP170719). This project was supported in part by the Genomic and RNA Profiling Core at Baylor College of Medicine with funding from the NIH S10 grant (1S10OD036427). The Baylor College of Medicine Single Cell Genomics Core is supported by NIH Shared Instrument Grants (P30CA125123, S10OD032189), and CPRIT grant RP200504. Work done by the 3D Organoid Core is supported by NIH U19 grant AI157981. We thank the core staff, in particular Devon Suske, Luqiong Wang, and Esmanur Tokcan, for their technical assistance.

## AUTHOR CONTRIBUTIONS

Conceptualization: JJZ, AWM, KAP; Methodology: JJZ, CMR, SEO, AK, ART, SEB; Investigation: JJZ, CMR, SEO; Formal analyses: JJZ, CMR, CC; Resources: RFC, AES; Funding acquisition: SEB, AWM, KAP; Writing – original draft: JJZ, KAP; Writing – review and editing: all authors.

