## Supplemental Figures 1-2 for "Human bladder organoids model urinary tract infection and bacteriophage therapy"

### Contents:

Supplemental Figures 1-2

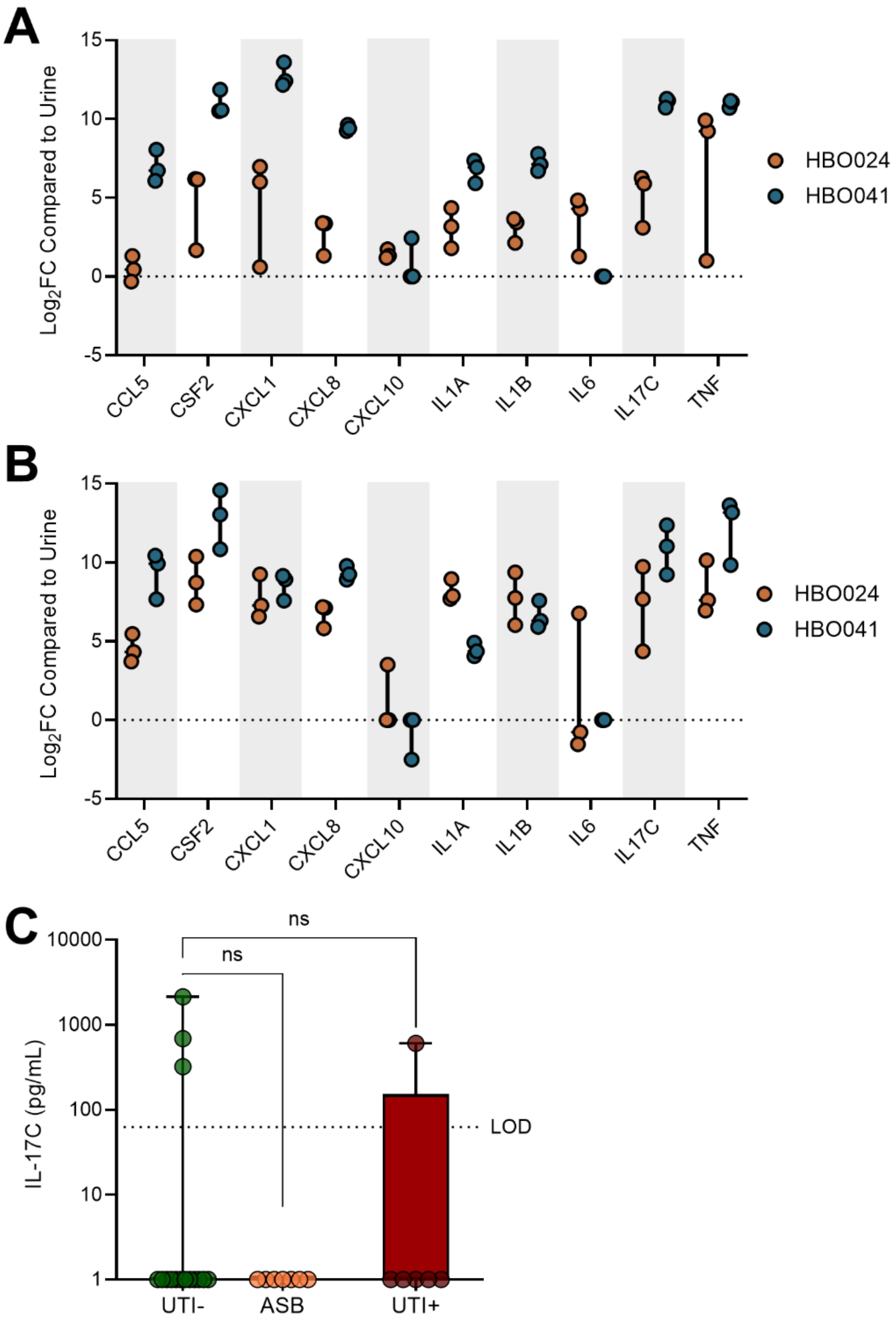


**Supplementary Figure 1: Comparison of relative cytokine expression between the two lines following UPEC infection.** $Log_2$ fold-change in inflammatory-related cytokines compared to urine control for both organoid lines after 4 (A) and 24 (B) hours of infection. This data is the expanded version of the heatmap shown in **Fig. 3E**. (C) IL-17C quantification from the urine of women with UTI, asymptomatic bacteriuria (ASB), or healthy controls. Data were analyzed using Kruskal-Wallis test with Dunn's multiple comparisons (C).

A

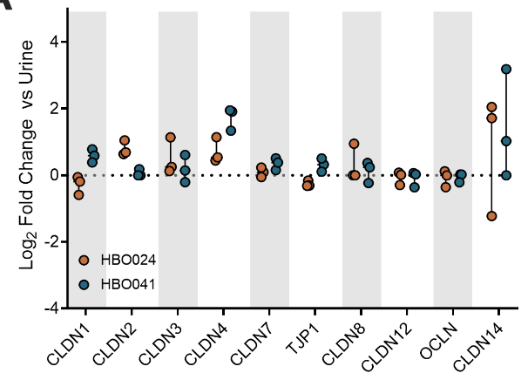


B

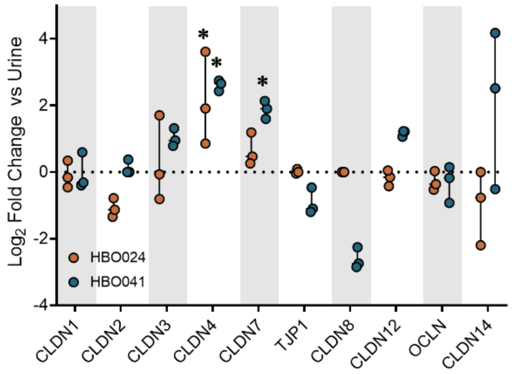


C

| | HBO024 4h | | | HBO041 4h | | | HBO024 24h | | | HBO041 24h | | |
|---|---|---|---|---|---|---|---|---|---|---|---|---|
| Gene | Log2FC | pvalue | padj | Log2FC | pvalue | padj | Log2FC | pvalue | padj | Log2FC | pvalue | padj |
| CLDN1 | -0.2376949 | 1 | 1 | 0.55922082 | 1 | 1 | -0.1070513 | 1 | 1 | -0.0765988 | 1 | 1 |
| CLDN2 | 0.87436107 | 1 | 1 | -0.2241595 | 1 | 1 | -1.0543127 | 1 | 1 | -0.0393984 | 1 | 1 |
| CLDN3 | 0.45725976 | 1 | 1 | 0.12745249 | 1 | 1 | 0.43268962 | 1 | 1 | 0.99844276 | 1 | 1 |
| CLDN4 | -0.0442139 | 1 | 1 | 1.69198336 | 0.06299358 | 1 | 2.39128321 | 0.00018538 | 0.00933724 | 2.61203607 | 1.48E-05 | 0.00029652 |
| CLDN7 | 0.08490005 | 1 | 1 | 0.309171 | 1 | 1 | 0.65289837 | 1 | 1 | 1.91104044 | 0.00114655 | 0.01302329 |
| CLDN8 | 0.73780259 | 1 | 1 | 0.22370675 | 1 | 1 | 0 | 1 | 1 | -2.6207435 | 0.04939415 | 0.26669336 |
| CLDN12 | -0.0803435 | 1 | 1 | -0.0826325 | 1 | 1 | -0.1566982 | 1 | 1 | 1.16991179 | 0.40310696 | 1 |
| CLDN14 | 1.57213326 | 0.81781232 | 1 | 4.32342076 | 0.19875585 | 1 | -2.2387161 | 0.63443286 | NA | 3.09548257 | 0.36814108 | NA |
| ZO-1 (TJP1) | -0.2519829 | 1 | 1 | 0.32084132 | 1 | 1 | 0.01596271 | 1 | 1 | -0.947094 | 1 | 1 |
| Occludin (OCLN) | -0.0499858 | 1 | 1 | -0.0334613 | 1 | 1 | -0.2690963 | 1 | 1 | -0.2590782 | 1 | 1 |

D

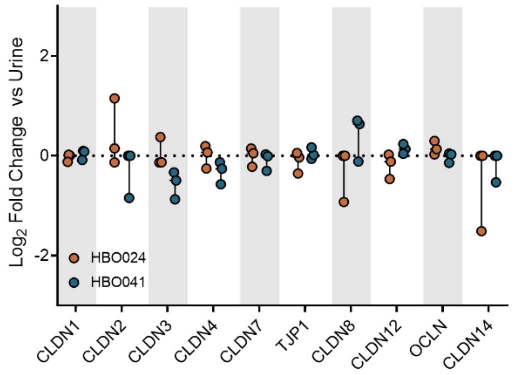


E

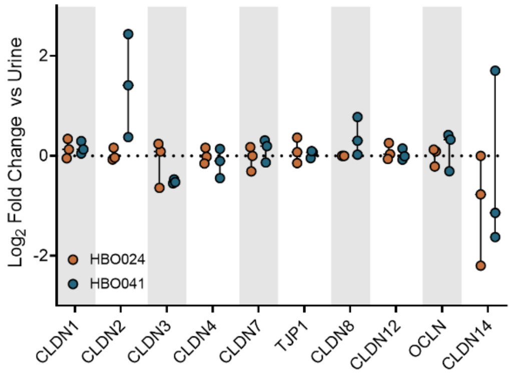


F

| | HBO024 4h | | | HBO041 4h | | | HBO024 24h | | | HBO041 24h | | |
|---|---|---|---|---|---|---|---|---|---|---|---|---|
| Gene | Log2FC | pvalue | padj | Log2FC | pvalue | padj | Log2FC | pvalue | padj | Log2FC | pvalue | padj |
| CLDN1 | -0.0284326 | 1 | 1 | 0.04743559 | 1 | 1 | 0.14210098 | 1 | 1 | 0.16593575 | 1 | 1 |
| CLDN2 | 0.25338701 | 1 | 1 | -2.180711 | 0.47764088 | 1 | 0.02325383 | 1 | 1 | 3.33246132 | 0.09030864 | 1 |
| CLDN3 | -0.0029352 | 1 | 1 | -0.4825235 | 1 | 1 | -0.122093 | 1 | 1 | -0.5174662 | 1 | 1 |
| CLDN4 | -0.0179751 | 1 | 1 | -0.272392 | 1 | 1 | -0.0156606 | 1 | 1 | -0.1035021 | 1 | 1 |
| CLDN7 | -0.0280815 | 1 | 1 | -0.1324321 | 1 | 1 | -0.0462708 | 1 | 1 | 0.11820304 | 1 | 1 |
| CLDN8 | -1.8856951 | 0.61467157 | 1 | 0.39751798 | 1 | 1 | 0 | 1 | 1 | 0.32974321 | 1 | 1 |
| CLDN12 | -0.1997877 | 1 | 1 | 0.14022655 | 1 | 1 | 0.08044955 | 1 | 1 | 0.03538591 | 1 | 1 |
| CLDN14 | -2.3878092 | 0.60720594 | 1 | 0.61089941 | 1 | 1 | -3.6525448 | 0.30857678 | 1 | -0.5115528 | 1 | 1 |
| ZO-1 (TJP1) | -0.1211667 | 1 | 1 | 0.01706251 | 1 | 1 | 0.11674433 | 1 | 1 | 0.03745133 | 1 | 1 |
| Occludin (OCLN) | 0.14058459 | 1 | 1 | -0.005417 | 1 | 1 | 0.00332819 | 1 | 1 | 0.17049534 | 1 | 1 |

**Supplementary Figure 2: Regulation of HBO tight junction proteins following UPEC infection and HP3 treatment.** Log2 fold-change in expression of bladder tight junction proteins following UPEC infection after (A) 4 and (B) 24 hours of UPEC infection, relative to urine only treated HBOs. (C) Table showing fold-change and p-values for bladder tight junction protein expression following UPEC infection. Log2 fold-change in expression of bladder tight junction proteins following HP3 treatment after (D) 4 and (E) 24 hours of incubation, relative to urine only treated HBOs. (F) Table showing fold-change and p-values for bladder tight junction protein expression following HP3 treatment.
